# Trypanocidal activity of the anthocyanidin delphinidin, a non-competitive inhibitor of arginine kinase

**DOI:** 10.1101/2020.06.23.167452

**Authors:** Edward Valera-Vera, Chantal Reigada, Melisa Sayé, Fabio A. Digirolamo, Facundo Galceran, Mariana R. Miranda, Claudio A. Pereira

**Author notes:** **Corresponding author:** C. Pereira, IDIM, Av. Combatientes de Malvinas 3150, (1427) Buenos Aires, Argentina.

## Abstract

The enzyme arginine kinase from *Trypanosoma cruzi* (TcAK) catalyzes the interconversion of arginine and phosphoarginine to maintain the ATP/ADP cell balance, and is involved in the parasites energetic homeostasis and stress responses. Using virtual screening approaches, some plant-derived polyphenolic pigments such as anthocyanidins, were predicted to inhibit TcAK activity. In this work, it was demonstrated that the anthocyanidin delphinidin showed a non-competitive inhibition mechanism of TcAK *in vitro* (Ki arginine = 1.32 μM and Ki ATP = 500 μM). Molecular docking simulations predicted that delphinidin occupies a hydrophobic pocket close to the ATP/ADP binding site. Delphinidin also exerted trypanocidal activity over *T. cruzi* trypomastigotes with a calculated IC_50_ of 19.51 μM. Anthocyanidins are low-toxicity natural products which can be exploited for the development of trypanocidal drugs with less secondary effects than those currently used for the treatment of Chagas disease.

---

The order Kinetoplastidae contains flagellated protozoan organisms which include many human pathogens from the genera *Trypanosoma* and *Leishmania. Trypanosoma cruzi* is the causative agent of Chagas disease, a parasitic zoonosis affecting approximately 7 million people in the Americas (https://www.who.int/news-room/fact-sheets/detail/chagas-disease-(american-trypanosomiasis)). Chagas disease is curable only if the treatment is applied during the early acute phase of the infection. However, in the chronic phase, antiparasitic drugs could only prevent the progression of the disease. About 30% of chronically infected patients present cardiac alterations and up to 10% develop a mixed form, involving cardiac and digestive symptoms, all of which require specific medical treatment^1^. Currently, there are only two medicines available for the treatment of Chagas disease, benznidazole and nifurtimox. Both drugs are effective in the acute phase but not in the chronic phase of the disease and have serious side effects. However, they could not be replaced or improved during the last 50 years^2^. This highlights the urgent need for more efficient drugs or alternative treatments, being one interesting possibility the search for bioactive compounds in the large universe of natural products.

Plants-derived natural products present a wide variety of biological and pharmacological properties. Anthocyanidins are flavonoid pigments that belong to a large family of polyphenols, and their glycosides are known as anthocyanins. These water-soluble compounds have large health benefit effects, most of them related to its antioxidant activity^3^. For example, anthocyanidins were active against night blindness, cancer, obesity and heart diseases^4–8^. In addition, various studies reported beneficial properties on bone health with the dietary intake of anthocyanins, such as delphinidin^4, 9^.

About thirty anthocyanidins were identified as the chemical core of the large group of anthocyanins but only six of them, including delphinidin, cyanidin, malvidin, pelargonidin, peonidin, and petunidin, are ubiquitous and of dietary importance^10^. In *T. cruzi*, the enzyme arginine kinase (AK, E.C. 2.7.3.3) is considered a potential drug target for Chagas disease. AK belongs to a wide family of guanidine phosphotransferases, including creatine kinase, and is involved in ATP buffering in cells with fluctuating energy requirements, catalyzing the interconversion between phosphoarginine and ATP^11^. Its potential as a drug target is supported mainly by the fact that AK is present in some invertebrates and trypanosomatid organisms but not in mammals^12, 13^. Additional evidence was provided in studies on *Trypanosoma brucei*, the etiological agent of the human African trypanosomiasis, in which AKs knockout is lethal under oxidative stress conditions^14^.

It was demonstrated that inhibition of *T. cruzi* AK (TcAK) by polyphenolic compounds present in green tea, specifically catechins, strongly affects the epimastigote cells viability^15, 16^. Taking into account these data, in a previous work we used an *in silico* approach, based on molecular docking techniques, in order to identify possible TcAK inhibitors in a database of antioxidant phenolic structures. Resveratrol, a natural stilbene, was chosen in that work for further *in vitro* assays due to its easy availability and low cost. However, resveratrol showed moderate effects not only as an AK inhibitor but also as an anti-*T. cruzi* agent^17^. Interestingly, in the same study the results of the TcAK molecular docking showed that the compounds interacting with the arginine binding site with lowest free binding energy values were three anthocyanidins: delphinidin, pelargonidin and cyanidin^17^. Therefore, delphinidin was chosen as a representative molecule to evaluate its potential not only as TcAK inhibitor but also as trypanocidal agent.

To validate that delphinidin is an inhibitor of TcAK and determine the inhibition mechanism, enzyme activity of the recombinant enzyme was measured^18^ varying arginine concentrations up to 0.4 mM with saturant concentrations of ATP (1.5 mM), and increasing delphinidin concentrations up to 2 μM. The resulting data was fitted to the Michaelis-Menten equation to calculate the kinetic parameters Km and Vmax (Fig. 1A and Supplementary Material), resulting in a 3.7 folds decrease of the Vmax (from 2.39 to 0.65 AU/sec) with increasing delphinidin concentrations (*p* = 0.0009), while Km showed no statistically significant difference between the different delphinidin concentrations (average = 0.70 mM; *p* = 0.4439). The decrease in Vmax excludes a competitive mode of inhibition, and the unvarying Km suggests a non-competitive mechanism with an inhibition constant Ki of 1.32 (± 0.11) μM. The same procedure was performed using ATP at different concentrations up to 1.6 mM and saturating concentrations of arginine (10 mM) (Fig. 1B and Supplementary Material). A decrease in the Vmax value of about 1.8 folds was observed (from 3.63 to 2.06 AU/sec) using delphinidin concentrations up to 0.3 mM (*p* = 0.0015) while Km values remain constant (average = 0.19 μM; *p* = 0.9649), indicating a non-competitive mechanism of inhibition regarding ATP, with a calculated Ki of 500 (± 79) μM.

**Figure 1.**
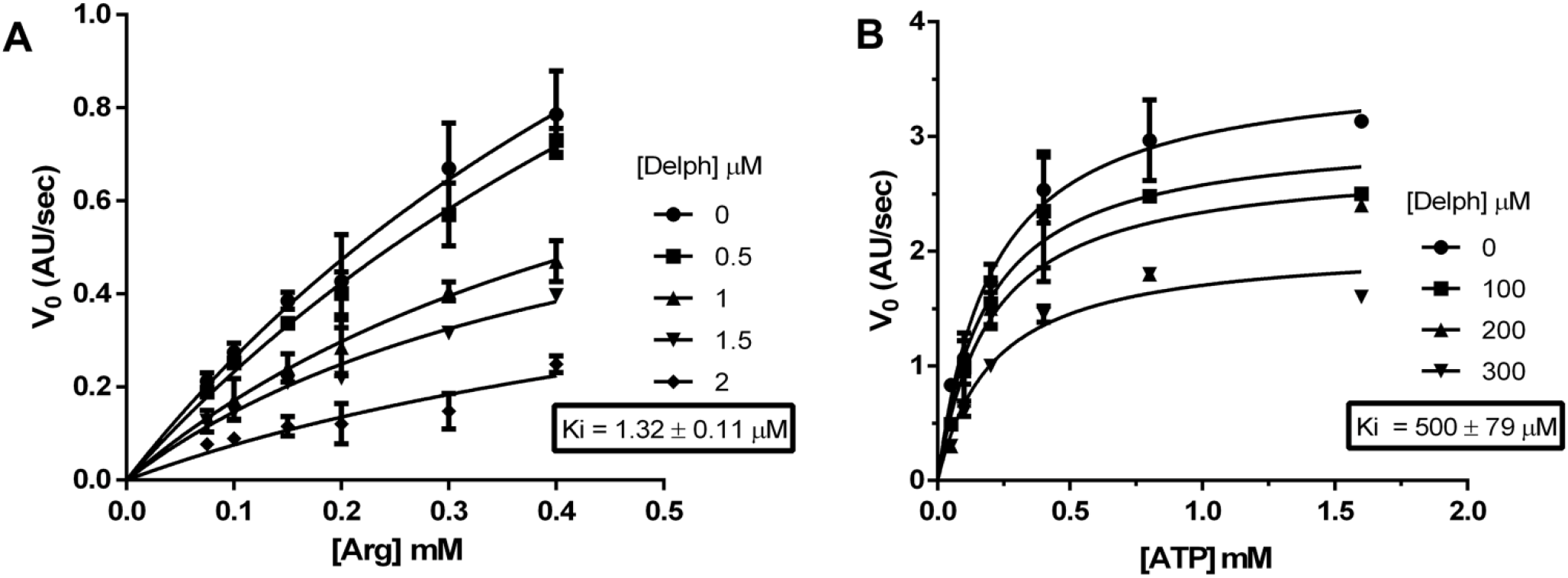
Kinetic analyses of TcAK inhibition. Enzyme activity of TcAK in presence of varying concentrations of arginine (A) or ATP (B) and delphinidin. Data was adjusted to the Michaelis-Menten equation to calculate kinetic parameters, Vmax and Km. Ki was obtained from adjusting the data to the non-competitive inhibition equation in GraphPad Prism. Complementary data to these graphics are detailed in Supplementary Material.

Based on the results obtained regarding the non-competitive inhibition kinetics of delphinidin for both substrates, it was decided to carry out a new and more detailed molecular docking analysis to identify possible interaction sites between delphinidin and TcAK other than the arginine binding one. First, we searched for the different pockets on the enzyme using the software fpocket 2.0 Two major pockets were identified, one comprising the ATP/ADP and arginine binding sites, and a second pocket on the enzyme surface (“exo-pocket”) (Fig. 2A). Molecular docking simulations were made on both pockets using AutoDock 4.5^20^ and AutoDock Vina^21^. Because scoring functions in docking programs are prone to error, the produced poses were re-scored using NNScore 2.0^22^, which is a neural network trained with experimental data in order to identify conformational features in high affinity binding modes. The two obtained poses for the exo-pocket that ranked best on the scoring algorithms had scores of −3.98 and −3.82 kcal/mol, being lower than the scores for the three best ranked poses on the substrate-binding pocket (−6.23, −6.15, and −5.89 kcal/mol). These data indicate that the most probable interaction site of delphinidin is the substrate-binding pocket. None of the poses docked on the active site occupied the arginine binding site, but were located close to the ATP/ADP binding site. One docking pose stood from the others (Fig. 2B), as both docking programs converged in a similar prediction that was also the most highly scored by the re-scoring function, mainly because of hydrophobic contributions. Visual examination highlighted the hydrophobic residues His185, Ile182, Trp221, and Ala312 as possible residues involved in the interaction with delphinidin.

**Figure 2.**
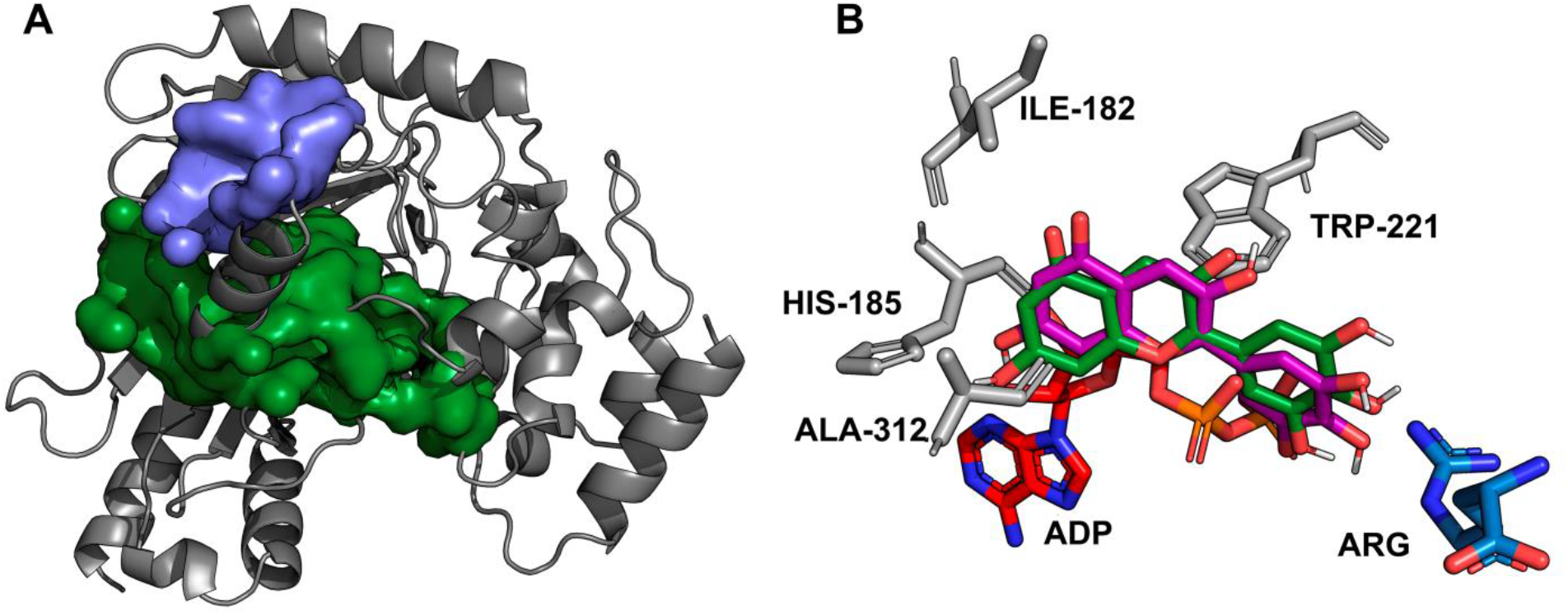
Molecular docking of TcAK and delphinidin. A) Molecular cavities in TcAK were predicted using fpocket 2.0. The pocket comprising the ATP/ADP and arginine binding sites is indicated in green, and the second one, surface or exo-pocket, is indicated in blue. B) Molecular structure of delphinidin and the converging docking poses from Autodock 4.5 and AutoDockVina, which obtained the best score according NNScore2.0. Figure also shows the experimental poses for ADP and arginine as well as the hydrophobic residues responsible of the predicted interaction (gray).

Since delphinidin showed a significant inhibition of TcAK activity, it was also tested for trypanocidal activity. The effect was evaluated on trypomastigotes because it is one of the therapeutically relevant stages of the *T. cruzi* life cycle. Culture derived trypomastigotes^23^ were incubated for 24 h with a delphinidin concentration range of 0 - 200 μM and the IC_50_ value (concentration of compound which gave 50% relative viability of parasites compared to the untreated control) was then determined. Delphinidin presented a trypanocidal activity with a calculated IC_50_ of 19.51 (± 1.05) μM (Fig. 3A). This value is similar to the obtained for benznidazole under the same experimental conditions; 14.9 (± 1.5) μM^24^. Vero cells exposed to delphinidin for 24 h in a concentration range from 0 to 200 μM, showed an IC_50_ of 39.16 (± 1.09) μM (Fig. 3B). These results demonstrate that delphinidin is 2-fold more specific over trypomastigotes than mammalian cells. All methods used in this work are detailed in the Supplementary Material.

**Figure 3.**
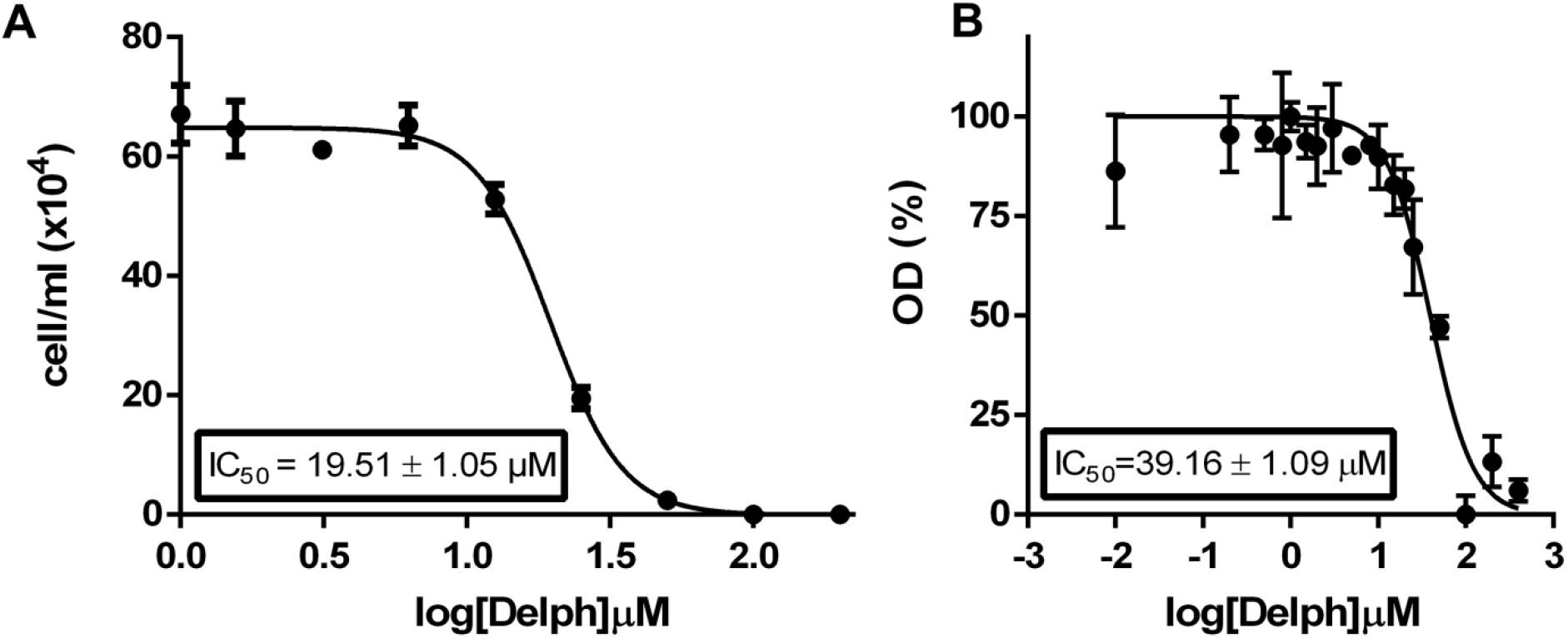
Trypanocidal effect of delphinidin. Dose-response curves of delphinidin over trypomastigotes of *T. cruzi* (A) and Vero cells (B), showing IC_50_ values obtained by non-linear regression. Treatments were performed during 24 h as indicated under “Materials and Methods”. OD (%) is the percentage of absorbance at λ = 570 nm of the crystal violet (vital staining) respect to the control without treatment.

The investigation of plants natural products as trypanocidal agents is an area of growing interest in the scientific community. A successful example is the work by Guida *et al*.^16^ reporting that the trypanocidal effect of catechins derived from green tea involves the inhibition of TcAK. Similarly, in the present work we studied the anthocyanidin delphinidin, which was previously postulated as TcAK inhibitor using virtual screening approaches^17^.

Delphinidin is a water soluble, low toxicity blue flower pigment, present in the genera *Viola* and *Delphinium*. It is also a natural pH indicator responsible for the blue-red color of different fruits such as grapes, cranberries, pomegranates, and bilberries^3^.

In contrast with previous results predicting that delphinidin interacts with the arginine binding site of TcAK^17^, in this work the obtained kinetic data shows that delphinidin does not compete with arginine when inhibiting TcAK as Vmax values decreased when delphinidine concentration was increased, while no statistically significant differences in Km values were observed according to a non-competitive inhibition model. The same is true when evaluating the inhibition mechanism of delphinidin against ATP, reason why new docking simulations were carried out comprising the totality of the substrate-binding pocket and a predicted exo-pocket.

Computational approaches suggest that binding of delphinidin might be occurring in a hydrophobic pocket close to the ATP/ADP binding site, even though inhibition kinetics show no competition between ATP and delphinidin. There are cases when a binding-site inhibitor shows a non-competitive behavior, for instance, this phenomenon have been observed in enzymes that experience many transition states to complete a catalytic cycle and return to a competent state that allows the start of a new cycle. This is the case for the *Limulus polyphemus* AK^25^. These enzymes have transition states where the inhibitor and not the substrate can bind yielding a non-competitive inhibition while both molecules bind to the same site^26^. Further evaluation of the inhibition mechanism in the reverse sense of the reaction may throw light on which transition state the inhibitor is binding.

Delphinidin showed anti-*T. cruzi* activity on trypomastigotes, one of the therapeutically relevant stages of the life cycle, with an IC_50_ value similar to that calculated for the reference drug, benznidazole^24^. In this sense, here we demonstrate the relevance of this molecule as a natural lead compound which can be exploited for the development of trypanocidal drugs with less toxicity than those currently used for the treatment of Chagas disease.

---

**Supplementary Table 1:** Kinetic parameters of TcAK inhibition

|  | Arginine |  |  |  |  | ATP |  |  |  |
| --- | --- | --- | --- | --- | --- | --- | --- | --- | --- |
| [Delph] μM | 0 | 0.5 | 1 | 1.5 | 2 | 0 | 100 | 200 | 300 |
| Vmax (app) AU | 2.39 ± 0.72 | 2.35 ± 0.47 | 1.15 ± 0.21 | 0.83 ± 0.14 | 0.65 ± 0.32 | 3.63 ± 0.15 | 3.05 ± 0.32 | 2.80 ± 0.45 | 2.06 ± 0.17 |
| Km (app) mM | 0.81 ± 0.33 | 0.91 ± 0.24 | 0.57 ± 0.15 | 0.47 ± 0.12 | 0.75 ± 0.52 | 0.20 ± 0.33 | 0.18 ± 0.06 | 0.19 ± 0.09 | 0.21 ± 0.17 |

Inhibition kinetics were performed by varying the concentrations of arginine between 0.075 and 0.40 mM while ATP stayed at a constant saturating concentration of 1.5 mM or varying ATP between 0.05 and 1.6 mM at a constant saturating concentration of arginine 10 mM. Delphinidin concentrations are indicated in the first row.

## Supplementary Methods

### Cells and parasites

Vero cells (African green monkey kidney) were cultured in MEM medium supplemented with 10% heat inactivated fetal calf serum (FCS), 0.15% (w/v) NaHCO_3_, 100 U/mL penicillin and 100 U/mL streptomycin at 37°C in 5% CO_2_ atmosphere. Trypomastigotes of the Y strain were obtained from the extracellular medium of Vero infected cells as previously described^1^.

### Trypanocidal activity assays

Trypanocidal activity in trypomastigotes was performed using 10^6^ cells/mL in 96-well plates which were incubated at 37°C for 24 h in the presence of different delphinidin concentrations. Trypomastigote number was determined by counting in a Neubauer chamber.

### Vero cell viability assay

Cytotoxicity against Vero cells was determined by the crystal violet staining assay. Briefly, the cells (10^4^ cells/well) were incubated in 96-well plates with delphinidin (or diluent only as a negative control) and maintained at 37°C for 24 h. At the end of the treatment, cells were fixed for 15 min, and stained with 0.5 % (w/v) crystal violet. After washing with water and drying, the staining was solubilized with 100 μl of methanol, and absorbance of the wells was measured at λ = 570 nm.

### Protein expression and purification

Briefly, the TcAK gene was amplified by PCR and cloned into the pRSET-A expression vector (Invitrogen). Protein expression was performed in *E. coli* strain BL21 (DE3). Recombinant TcAK was purified by affinity chromatography using a Ni-NTA Superflow resin (QIAGEN) and used for biochemical assays^2^.

### Arginine kinase activity and inhibition assays

Enzyme activity was measured by a spectrophotometric coupled-enzymes method^3^. The reaction mix consisted of 100 mM Tris-HCl buffer pH 8.2, 1.5 mM MgCl_2_, 0.5 mM DTT, 1.5 mM phosphoenol-pyruvic acid, 0.3 mM NADH, 5 units of lactate dehydrogenase (Sigma Chemical Company) and 5 units of pyruvate kinase (Sigma Chemical Company). The enzyme source was affinity purified recombinant TcAK. Inhibition kinetics were made by varying the concentrations of arginine (0.075, 0.10, 0.15, 0.20, 0.30, and 0.40 mM) while ATP stayed at a constant saturating concentration of 1.5 mM, with different delphinidin concentrations (0, 0.5, 1.0, and 2.0 μM). The same assay was also performed varying ATP concentrations (0.05, 0.1, 0.2, 0.4, 0.8, and 1.6 mM) with arginine 10 mM, a constant saturating concentration, and different concentrations of delphinidin (0, 0.1, 0.2, and 0.3 mM). Reaction was started by addition of the constant substrate. NADH oxidation was measured using a spectrophotometer at λ = 340 nm. Controls were performed using delphinidin in the presence of 1.5 mM ADP, to bypass the AK reaction, and the activity of the coupled enzymes was not affected. In addition, the measured absorbance of delphinidin at λ = 340 nm was negligible.

### Pocket prediction and molecular docking

Since the only experimentally determined structure TcAK on the Protein Data Bank (PDB: 2J1Q) lacks some critical residues (289 to 295 and 310 to 320), a homology model was produced by iterative template-based fragment assembly using the I-TASSER server^4^. Protein pockets on the model were predicted using the software fpocket 2.0^5^. 3D structure of delphinidin was downloaded from ZINC database (http://zinc15.docking.org/; ID: ZINC3777403). Preparation of the PDBQT file was performed using AutoDock Tools 1.5.6 6. The grid parameter file was generated with Autogrid 4.2.6 so as to surround the predicted pockets. AutoDock 4.5 was used for calculation of optimal energy conformations for the ligands interacting with the protein pockets running the Lamarckian Genetic Algorithm 100 times, with a population size of 300, and 2.7×10^4^ as maximum number of generations followed by conformational clustering of the obtained poses into groups with RMSD <0.2 nm. Complementary, the AutoDock Vina software^7^ was also used. All the obtained docking poses were visually analyzed using PyMOL (https://pymol.org/) and re-scored using the neural-network-based scoring software NNScorev2.0^8^.

### Statistics and data analysis

IC_50_ values were obtained by non-linear regression of dose-response logistic functions, using GraphPad Prism 6.01 for Windows. All experiments were performed in triplicate and the data are presented as mean ± standard deviation (SD). Vmax and Km values were compared using the Kruskal-Wallis test.

